# Isolating objective and subjective filling-in using the Drift Diffusion Model

**DOI:** 10.1101/2023.02.01.526391

**Authors:** Ron Dekel, Dov Sagi, Ativ Zomet, Dennis M. Levi, Uri Polat

## Abstract

Spatial context is known to influence the behavioral sensitivity (*d*’) and the decision criterion (c) for detection of low-contrast targets. Of interest here is the effect on the decision criterion. Polat and Sagi (2007) demonstrated that for a Gabor target positioned between two similar co-aligned high contrast flankers, observers’ reports of seeing the target (Hit and False-Alarm) decreased with increasing target-flankers distance. This effect was more pronounced when distance was randomized within testing blocks compared to when it was fixed. According to Signal-Detection-Theory (SDT), the latter result suggests that the decision-criterion is adjusted to a specific, distance-dependent combination of signal (S) and Noise (N) when the S and N statistics are fixed, but not when they vary across trials. However, SDT cannot differentiate between changes in the decision-bias (criterion-shift) and changes introduced by variations in S and N (signal-shift). To circumvent this limitation of SDT, we analyze reaction-time data (RT) within the framework of the Drift-Diffusion-Model (DDM). We performed an RT analysis of target-flanker interactions using data from Polat & Sagi (2007) and Zomet et al. (2008; 2016). The analysis revealed a stronger dependence on flankers for faster RTs and a weaker dependence for slower RTs. The results are explained by DDM, where an evidence accumulation process depends on the flankers via a change in the rate of the evidence (signal-shift), and on observers’ prior via a change in the starting point (criterion-shift), leading to RT-independent and RT-dependent effects, respectively. The RT-independent distance-dependent response-bias is attributed to the observers’ inability to learn multiple internal distributions required to accommodate the distance-dependent effects of the flankers on both the Signal and Noise.

## Introduction

Detection of an oriented target improves in the presence of similar, co-aligned, high contrast flankers (Morgan & Dresp, 1995; Polat & Sagi, 1993, 1994; Solomon & Morgan, 2000; Woods et al., 2002). For oriented Gabor targets, contrast sensitivity is doubled when the distance between the target and flankers is about three times the Gabor wavelength (Polat & Sagi, 1993). These spatial interactions are suggested to be a manifestation of brain processes involved in contour filling-in, in texture segmentation, and in perceptual grouping (i.e. contour integration) (Sagi, 1995; Zhaoping & Jingling, 2008). The earlier experiments, cited above, used the bias-free 2-Altrnative-Forced-Choice (2AFC), considered efficient in estimating visual sensitivity (d’), but they provide no insight into the perceived quality of targets, which is expected to be affected by filling-in processes (Anstis, 2010). Polat and Sagi (2007), employing the Yes/No method, found, in addition to detection facilitation, a distance dependent detection bias; observers’ tendency to report “target present” increased at short target-flankers distances regardless of target’s presence (Hit) or absence (False Alarms: FA). This suggests that the gap between flankers is filled in with task-relevant information, supporting the ‘filling-in’ hypothesis. Both the increased target sensitivity and the observed detection bias are thought to be caused by lateral interactions in the visual cortex, activated by the flankers. Report biases are also affected by decision strategies, possibly related here to statistical priors derived from the known characteristics of natural images (Geisler et al., 2001). To better understand the contributions of lateral-interactions and decision-strategies to the detection bias, we present a reaction time (RT) analysis of experimental results collected in the previous ‘Yes/No’ experiments (Polat & Sagi, 2007; Zomet et al., 2008; Zomet et al., 2016). The data was modelled using Signal Detection Theory (SDT) (Green & Swets, 1966) and the Drift Diffusion Model (DDM) (Ratcliff & McKoon, 2008; Ratcliff et al., 2016; Shadlen & Kiani, 2013). In the subsequent sections we elucidate the unified SDT-DDM framework employed to model the data.

### The SDT approach

Following SDT, it is assumed that observers base their decision on noisy sensory activity within the brain (referred to as the “internal response”), monotonically increasing with stimulus strength. For low contrast targets, the internal target-response distribution (referred to as Signal, *p*_*S*_*(x)*) may overlap with the internal noise distribution (representing no target, termed Noise, p_N_(x)), thus leading to detection errors (Figure 1). Consequently, ‘Yes’ responses can be correct (Hit, P_Hit_ = area under the green shaded curve in Figure 1A) or incorrect (False Alarm, P_FA_ = area under the red shaded curve in Figure 1A).

**Figure 1:**
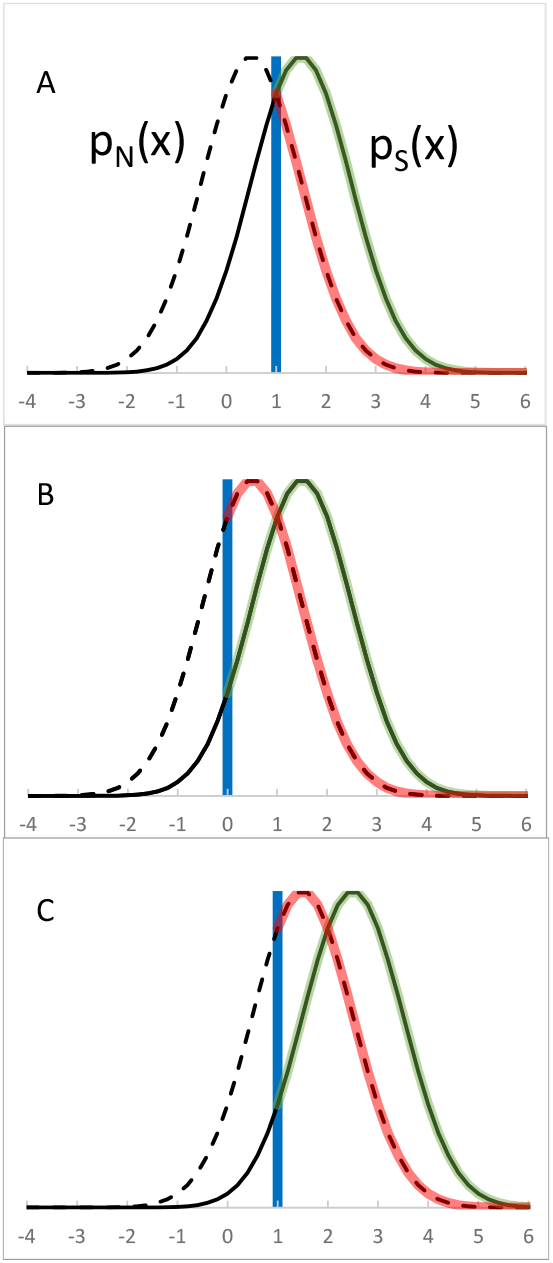
Illustration of the problem addressed in this work: **(A)** The standard SDT scheme, p_N_(x) the noise distribution (broken curve), p_S_(x) the Signal distribution (continuous curve), decision criterion (blue vertical line). Regions of Hit and FA are highlighted by green and red colors, respectively, with the area under the highlighted segments being P_HIT_ and P_FA_. **(B)** The decision criterion is shifted toward lower internal responses, thus increasing P_HIT_ and P_FA_. **(C)** p_N_(x) and p_S_(x) are shifted toward higher activity levels without a criterion shift relative to **(A)**, producing the same increase in P_HIT_ and P_FA_ as in **(B)**. Our goal here is to decide between B and C.

SDT provides tools to compute a decision criterion from the P_Hit_ and P_FA_, which is the normalized internal response level above which the observer produces a ‘Yes’ decision (depicted as the blue vertical line in Figure 1A). This criterion is assumed to be observer dependent, and can shift per task demands and available stimulus properties. However, when observing a specific change in Hit and FA rates, such as the increased rates seen in our experiments, SDT cannot distinguish between two potential causes: (1) a shift in the decision criterion toward lower response levels (Figure 1B), or (2) an elevation in activity levels at the target location, shifting both Signal and Noise distributions to higher response levels (Figure 1C). In the realm of SDT, the first cause - a criterion shift - is believed to depend on observers’ decision strategy, which is flexible and aims to optimize task outcome, including error rates, cost and value making it inherently ‘subjective’. On the other hand, the second cause is deemed sensory-driven, ‘objective’, and tied to the stimulus (such as flankers), influencing the system’s response. Regarding the experimental paradigm studied here, one might anticipate an increase in ‘Yes’ responses in the presence of flankers due to (1) observers’ expectation for the gap between flankers to be filled-in, or due to (2) increased sensory activity at the target location (as in Figure 1C) prompted by the input from the flankers. Both causes might reflect the visual system’s adaptation to the statistics of edge co-occurrence in natural images (Geisler et al., 2001).

### Linking SDT to DDM

To disentangle the inherent ambiguity within SDT and distinguish between the contributions driven by sensory activity and those influenced by decision criteria, we examined the experimental results using the framework of the DDM (Ratcliff & McKoon, 2008; Ratcliff et al., 2016; Shadlen & Kiani, 2013). Unlike SDT, DDM introduces a temporal dimension to the decision process, allowing RT based predictions. According to DDM, observers accumulate evidence both in favor of and against the presence of a target. Early models (Gold & Shadlen, 2001; Link, 1975; Stone, 1960) followed Wald’s (1947) Sequential Probability Ratio Test (SPRT), suggesting that each time interval produces a log-likelihood ratio (*LLR*) value. This value assesses the odds ratio for one stimulus being present versus the other, and it accumulates over time intervals until a decision is triggered. This happens when the accumulated value reaches one of two thresholds (bounds), e.g., +*a* and –*a*, for positive and negative decisions, respectively. The starting point of the accumulator (*sp*) can be selected to incorporate expectations, prior information and the subjective value of the decision, such as payoff and reward. The rate of evidence accumulation, termed “drift rate”, increases with the target sensitivity, resulting in faster attainment of the decision bounds. When the internal response offers no evidence for target presence or absence (for instance, when *p*_*S*_*(x)=p*_*N*_*(x)*), the drift rate (v) is zero. Positive and negative drift rates correspond to target-present and target-absent trials, respectively. Thus, within the SDT framework, we assume that the log likelihood ratio *LLR(x)=log(p*_*S*_*(x)/p*_*N*_*(x))* is integrated over time, where *p*_*S*_*(x)* and *p*_*N*_*(x)* (as illustrated in Figure 1) represent the momentary distributions of the sensory evidence (x) in the signal (S) and noise (N) trials. More formally, for a time-varying response *x(t)*, and accumulated value *L*, we have *L(t>0) = L(t-1) + LLR (x(t))*, with *L(0) = sp*. The mean drift rate (*v*) in *S* and *N* trials, *v*_*S*_ and *v*_*N*_ respectively, is assumed to be proportional to the expected value of *LLR(x(t))* over the corresponding *S* and *N* trials. A decision is reached when *L(t)≥a* (a positive decision) or when *L(t)≤-a* (a negative decision). Importantly, note that the effect of *L(0*) on *L(t)* is expected to diminish with time as *L(t)* accumulates evidence and noise.

While offering an efficient method for sequential hypothesis testing, a critical limitation of the likelihood model lies in its requirement for knowledge of the Signal and Noise distributions (necessary for each *x(t)* so that *LLR(x(t))* can be accumulated). This necessity is often deemed challenging, if not unattainable, particularly in typical psychophysical experiments characterized by a limited number of trials. An alternative approach, employed by DDM, directly integrates the sensory evidence (Ratcliff & McKoon, 2008; Shadlen & Kiani, 2013). Gold and Shadlen (2001) proposed the difference between the momentary response and the criterion level as an alternative to the likelihood ratio computation, though the method of criterion setting is left open. The approach presented here explicitly assumes, as described below, that in uncertain environments where observers encounter diverse stimuli with varying internal distributions, they fail to accurately estimate these distributions. Consequently, they base their decision on a mixture distribution, applying a single decision criterion to all stimuli (Gorea et al., 2005; Gorea & Sagi, 2000).

#### Interacting decision criteria

Consider the case where various stimuli are presented in an experiment, yielding stimulus dependent S and N distributions, as when varying target-flankers distance between trials (‘Mix’ condition, Figure 2). The findings of Gorea and Sagi (2000) suggest that observers are unable learn the individual distributions, as required for optimal performance. Instead, they merge all S and N distributions (related to the different distances) into single S and N distributions, estimated to represent the average of the individual distributions. For the decision-making process, observers employ only one criterion that is optimized for these single S and N distributions. Consequently, it is predicted that only one accumulator is used in the mixed condition, with the estimated evidence for/against target presence being blind to the originating, distance dependent, distribution. Therefore, we expect zero evidence to correspond to the presence/absence of targets regardless of the specific flanker configurations. In essence, this is determined globally by amalgamating the diverse distributions related to the various target-flanker distances (as depicted in Figure 3).

**Figure 2:**
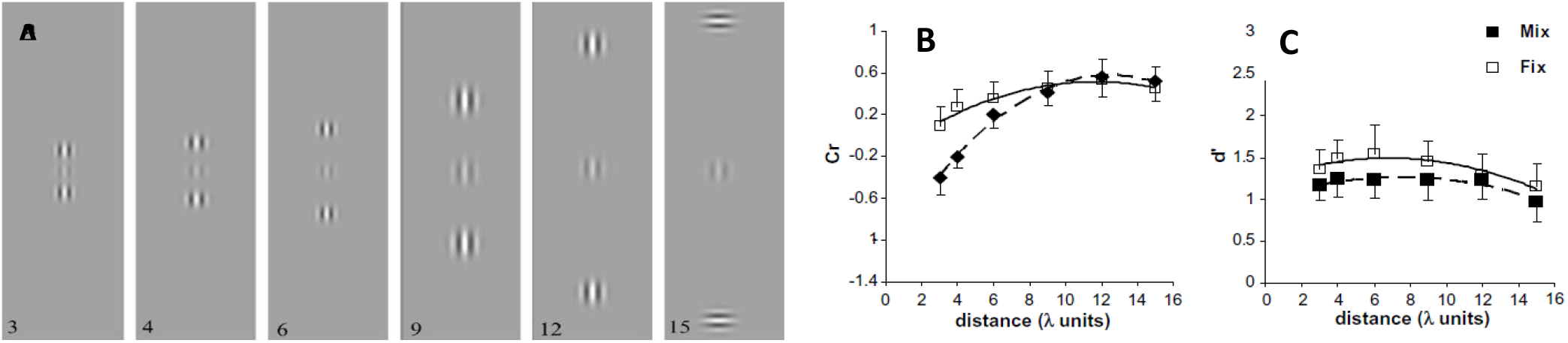
Lateral masking and the SDT criterion. **(A)** Stimuli used in Polat & Sagi (2007). Observers detected the presence vs. the absence of the low-contrast target stimulus presented between two high-contrast flankers aligned with the target (except for the 15λ condition used for reference). The distance between the target and flankers was varied (3, 4, 6, 9, 12, and 15λ) between blocks of trials (the Fixed condition) or within one block of mixed trials (the Mixed condition). (**B and C**) Group results from Polat & Sagi (2007, Figure 3): the effect of flankers on target detection, as measured by the signal detection theory (SDT) (**B**) criterion (Cr), and (**C**) sensitivity (d’). Shown are the means ± SEM across 7 observers (see the Methods). Note the close to uniform criterion level in the Fix condition compared with the larger range in the Mixed condition. d’ does not differ much between conditions, though it is somewhat lower in the Mix condition, possibly due to the increased uncertainty involved in this condition. The d’ curves show lateral facilitation at shorter distances, though smaller than the typical facilitation observed with the standard 2AFC method (Polat & Sagi, 1993; Polat & Sagi, 2007).

**Figure 3:**
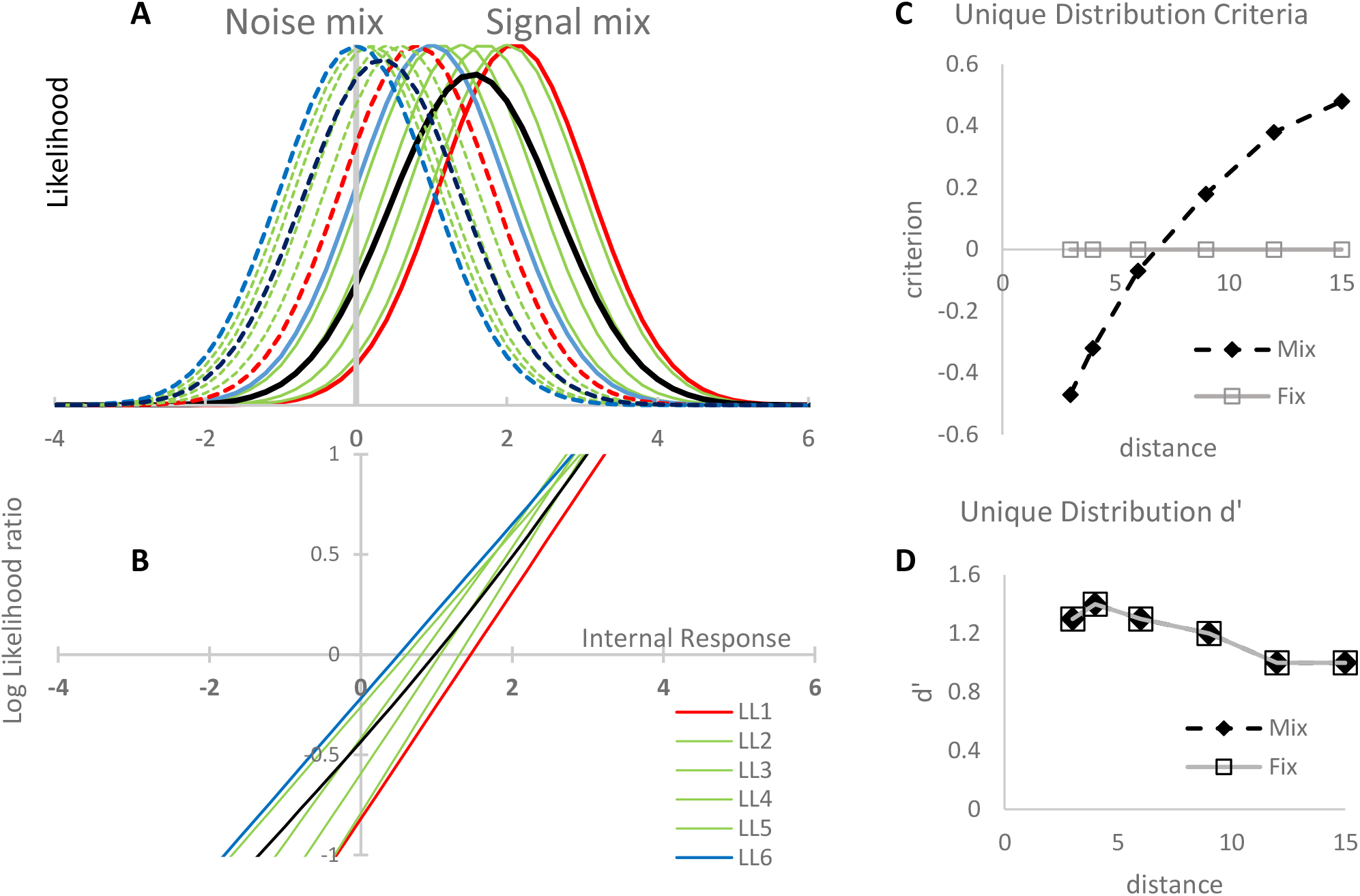
The proposed SDT modeling of the measured criterion change. (A) Internal response distributions thought to be involved in the studied task (normal distributions, σ=1). There are 6 noise distributions (broken lines) and 6 signal distributions (continuous lines), with the shortest and longest target-flanker distances denoted in red and blue, respectively. We assume that the observers cannot estimate all of these distributions when encountered randomly within a block of trials, and instead use the mean distributions of Noise and Signal (the broken and continuous black curves, respectively). (B) Log likelihood ratio (LLR) functions computed for Noise and Signal pairs in A. We expect a ‘Yes’ response when LLR>0 and ‘No’ otherwise. Observers are assumed to base their decisions on the black curve (LLR of the means) instead of the specific distance curves; thus, they are biased. (C) Predicted criterion shifts derived from B. For the Fix condition, we assume ideal observers (unbiased). (D) d’ values associated with the distribution pairs are presented in A. These values are computed as the difference between the paired Signal and Noise distributions.

In this study, we undertook an analysis of the task involving detection of a low-contrast Gabor patch in the presence of flankers (Figure 2A). The effect of the flankers depends on the target-flanker distance, showing improved sensitivity at intermediate target-flanker distances (2λ < distance < 8λ) (where λ is the wavelength of the Gabor stimuli). This phenomenon, known as range-dependent lateral facilitation, induces an impression of filling-in, leading to an elevation in the False Alarm (FA) rate in Yes/No detection experiments (Polat & Sagi, 2007). This effect can be quantified by the decision criterion (Figure 2B) as defined in SDT. While the criterion can be adjusted, though not perfectly, by observers when different target-flanker distances are blocked (Figure 2B, ‘Fix’), it was found to depend on the flankers when the different target-flanker distances are mixed (Figure 2B, ‘Mix’). This result can be explained by observers failing to independently adjust multiple criteria when stimulus strength, that is distance dependent, is randomized within a block of trials, as illustrated in Figure 3. To explain these effects, the flankers can be assumed to increase the response level of the target stimulus population, with the effect depending on the target-flanker distance. Figure 3 illustrates the challenge faced by observers in such a task. In each trial, an internal response is drawn from one of twelve distributions, corresponding to six distances, both with and without the target present (6×2, as shown in Figure 3A). For an observer to make unbiased decisions, these twelve distributions need to be correctly estimated, with an unbiased criterion set for each distance based on the corresponding pair of distributions (Noise and Signal). Instead, we propose that observers utilize only two distributions - the average Noise distribution and the average Signal distribution (as depicted the black curves in Figure 3A). The ratio between these distributions provides the input to their decision (Yes if *LLR(x)>0*, No otherwise, ‘x’ being the internal response, see the black curve in Figure 3B). In a blocked condition (where distance is Fixed), observers can possibly estimate the specific distance-dependent distribution and derive an unbiased likelihood ratio value for each distance (illustrated by the colored curves in Figure 3B). The anticipated decision criteria for the example outlined in Figure 3A are depicted in Figure 3C.

It is also possible that observers bias their decision for each distance independently, based on prior knowledge about the statistics of filling-in between contour elements, as shown by Gorea and Sagi (2000) for targets having different prior probabilities. Such biases may affect the predictions shown in Figure 3C, depending on the specific priors assigned to different distances. Thus, as outlined earlier, we can anticipate the presence of two contributing sources to the criterion effect: (1) activity based, affecting the statistics of internal responses (leading to shifted *p*_*S*_*(x)* and *p*_*N*_*(x)* distributions, akin to Figure 1C), and (2) biases corresponding to observers’ prior knowledge of the stimuli and the task at hand (resulting in a shifted criterion, similar to Figure 1B).

Note that the above discussion concerns the mechanisms of criterion setting and is mute regarding observers’ sensitivity to changes in target contrast. Sensitivity depends on the internal response gain, which corresponds to the difference between the means of the S and N distributions. In the context of SDT, sensitivity is described by d’ that is computed in a way that is assumed to be criterion independent (Figure 2C, Figure 3D).

#### A criterion that disappears with RT and that does not disappear with RT

In prior studies involving tasks like detection, Tilt After Effect, and Tilt Illusion, we have demonstrated that perceptual biases arising from shifts in decision criteria (as in Figure 1B) diminish as reaction times (RTs) lengthen. In contrast, biases attributed to sensory interactions (as in Figure 1C) remain unaffected by RT (Dekel & Sagi, 2020b). DDM provides a coherent explanation for these phenomena. In the DDM framework, shifts in decision criterion are implemented by modifying the starting point of the evidence accumulation process. Consequently, when the process takes longer to terminate (manifesting as slower RT), decisions are less biased due to noise accumulation (Dekel & Sagi, 2020b). Conversely, alterations in decision criteria arising from sensory interactions are characterized by a change in the rate at which evidence accumulates. As a result, such effects exhibit minimal dependence on RT. In the present work, we adopt this approach to analyze lateral masking data collected from previous studies, which had not been previously subjected to RT analysis (Polat & Sagi, 2007; Zomet et al., 2008; Zomet et al., 2016).

#### The present project

The experimental findings presented above show a clear dependency of decision bias on the target-flanker distance when the different distances are mixed but not so much when observers are presented with a fixed distance. The mixture distribution model presented above predicts distance dependent decision biases (i.e. criterion shifts) when distances are mixed, caused by internal-response shifts interfering with the formation of efficient representations of the internal distributions associated with the different stimuli. In addition, there may be expectation dependent decision biases resulting from observers adopting different decision rules for different stimuli. These expectation based biases are presumed to disappear at slow RTs, while biases predicted by the mixture distribution model are expected to be present at slower RTs in experimental conditions where different stimuli (here different distances) are mixed but not when distance is fixed. Based on these arguments we present four predictions:

P1: The criterion dependence on distance (Figure 1B) is expected to be larger at faster RTs, but still present at slower RTs when trials of different distances are mixed but not when the distances are blocked.

P2: The criterion dependence on distance is expected to be larger at faster RTs due to biases introduced by the starting point of the accumulator (*sp*=*L(0)*). These biases are reduced with increasing RT due to accumulation of internal noise. However, high levels of external noise (Zomet et al., 2016), introduced at *t=0*, are expected to dominate the internal noise and to reduce the effect of the accumulation starting point. Thus, we predict RT to have reduced effect on the criterion dependence on distance.

P3: The criterion dependence on distance is expected to decrease with decreasing slope of the log likelihood function presented in Figure 3B. It becomes evident that the slope decreases when the *S* and *N* distribution width (σ) is increased. For the specific model presented, assuming normal distributions with constant *S* and *N* means, the slope is proportional to *(‹S›-‹N›)/σ*^*2*^. Thus, the criterion dependency on distance is expected to decrease with increasing σ, and to vanish at slower RTs in the presence of high noise. For the case where the *S* and *N* difference is increased with σ, the dependency of criterion on distance is expected to be preserved at higher noise levels. These predictions are tested by analyzing results from experiments where external noise is added to the target (Zomet et al., 2016).

P4: The criterion dependence on distance is expected to be larger at faster RTs, thus to be reduced in slower observers (as discussed above, according to DDM, the starting-point dependent bias decreases with RT (Dekel & Sagi, 2020a)). Here we analyze data from a group of observers diagnosed for depression (Zomet et al., 2008), showing slower RTs. We expect the faster RTs of this group to have reduced dependence of criterion on distance.

In the following we test these predictions.

## Methods

### Experimental data

Here we analyzed unpublished RT data from previously reported experiments (Polat & Sagi, 2007; Zomet et al., 2008; Zomet et al., 2016), as detailed below and summarized in Table 1. All experiments measure the detection of low-contrast vertical Gabor patch “targets” in the presence of two lateral high-contrast Gabor patch “flankers” (Figure 2A). Flankers were located at varying distances from the target (3-15 λ where λ is the wavelength of the Gabor patch). In all experiments, the 15λ distance used horizontally oriented flankers, presumably nulling any lateral interaction with the vertical target, whereas all other distances used vertically oriented targets. Auditory feedback was used to mark detection errors. From Polat and Sagi (2007), we recovered the original data for most observers (six and five observers out of seven, for ‘Mix’ and ‘Fix’, respectively). In the remaining publications, all the original data were recovered, as well as an unpublished pilot study employing external noise, as in (Zomet et al., 2016), with numerous repetitions per participant. The observers in all experiments differed, with the exception of those in Polat and Sagi (2007), where five observers were shared between ‘Fix’ and ‘Mix’. The stimulus parameters and experimental setup, detailed below, were nearly identical in the different experiments (see the differences in Table 1).

**Table 1.** Summary of the experimental data.

| Publication | Experiment | Design |  |  | Target Contrast | Participants |  |  |  |
| --- | --- | --- | --- | --- | --- | --- | --- | --- | --- |
| | | Target-flanker distance ($\lambda$) | External noise levels | Mixing of target-flanker distances | | N | Gender (M/F) | Age (Mean ± SD) | #Trials per participant (Mean ± SD) |
| Polat & Sagi (1993) | Mix | 3,4,6,9,12,15* | - | Mixed | Fixed per observer | 6** | 5/1 | 17 ± 2 | 3550 ± 1400 |
| | Fix | 3,4,6,9,12,15* | - | Blocked | Fixed per observer | 5** | 4/1 | 17 ± 2 | 5400 ± 2800 |
| Zomet et al. (2008) | Controls | 3,4,6,9,12,15* | - | Mixed | Fixed per observer | 32 | 14/18 | 42 ± 15 | 110 ± 130 |
| | Depress | 3,4,6,9,12,15* | - | Mixed | Fixed per observer | 27 | 15/12 | 48 ± 15 | 120 ± 190 |
| Zomet et al. (2016) | Main experiment | 3,4,6,15* | 0,0.5,1,2 | Mixed | Fixed per observer | 12 | 2/10 | 24 ± 3 | 930 ± 50 |
| | Unpublished pilot | 3,4,6,9,12,15* | 0,0.5,0.75, 1,1.5,2 | Mixed | Fixed per observer, increasing with noise | 7 | 2/5 | 26 ± 4 | 2100 ± 50 |

\* The 15λ distance uses flankers with a horizontal orientation.
\*\* Missing data from the original study.

### Stimuli and Procedure (Polat & Sagi, 2007)

The stimuli consisted of Gabor patches with wavelength λ=0.11º, modulated from a background luminance of 40 cd.m^-2^ (Figure 2). Stimuli were presented on a Philips multiscan 107P color monitor, using a PC system. The effective size of the monitor screen was 24 × 32 cm, which at the used viewing distance of 150 cm subtends a visual angle of 9.2 × 12.2 degrees. Observers’ viewed the stimuli binocularly in a dark cubicle, where the only ambient light came from the display screen.

Stimuli consisted of a low-contrast Gabor-target and two high-contrast (60%) Gabor-flankers (Figure 2). Target-flanker distances used were 1, 2, 3, 4, 6, 9, 12 and 15λ. Baseline thresholds, against which spatial interactions were compared, were obtained using orthogonal target and flankers with an inter-element distance of 15λ. The contrast of the target was constant for each observer at all target-flanker distances with the exact value depending on the contrast detection threshold for that observer (range 3-4%). Using the Yes/No task, the observers were asked to detect the target which was shown in a single presentation, and to report whether the target was present (Yes) or absent (No) by pressing the left and right mouse keys, respectively. A visible fixation circle indicated the location of the target, it disappeared when the trial was started. Observers activated the presentation of the trials at their own pace. When the 2AFC task was used, there were two stimulus intervals (80ms each), presented 800ms apart, both containing the flankers but with the target presented in only one of the intervals.

There were two main experimental procedures. In the “Mix” procedure, the trials with different target-flanker distances were presented in random order, while in the “Fix” procedure the different target-flankers distances were blocked. In the Fix procedure, target-flanker distance was change randomly between blocks. In both procedures, each distance was presented 50 or 20 (depending on the experiment) times in a session with target present on about half of the trials (probability of 0.5).

### Stimuli and Procedure (Zomet et al., 2008)

The stimuli and task were the same as in Polat and Sagi (2007), using the ‘Mix’ procedure, but with a wavelength λ=0.16º and stimulus duration of 100ms. Stimuli were presented on a Viewsonic (Brea, California) E70 color monitor with display dimensions as above. The two groups (patients and controls) did not differ significantly (*p* = .097) in contrast threshold. The mean contrast threshold of the control group was 5.12, and that of the patient group was 6. In each session, 20 trials for each target-flanker separation were presented, with a total of 120 trials per session.

The patients were hospitalized in the Psychiatry Department at Sheba Medical Center. These patients were diagnosed by psychiatrists as suffering from Major Depression Disorder (MDD), according to DSM-IV. All patients were found to be currently depressed during our testing period and were being treated by antidepressants and benzodiazepine medications (see Zomet et al (2008) for more details).

### Stimuli and Procedure (Zomet et al., 2016)

The method was similar to the method described above (Zomet et al., 2008), but with only 4 target-flanker distances (3, 4, 6, 15 lambda, (λ)) in the main experiment. Stimuli were displayed on a Sony multiscan G400 color monitor (1024 × 768 pixels). The effective size of the monitor was 26 × 35 cm, which at a viewing distance of 150 cm subtended a visual angle of 9.7 × 11.4 degrees.

White noise (containing a broad range of random orientations and spatial frequencies) at differing levels of contrast was presented at the target location and was superimposed on the target when present. For each observer, the noise contrast was normalized to his/her noise threshold detection threshold, measured separately estimated using an adaptive staircase method (79% correct), and was presented at 0, 0.5, 1, and 2 times their noise detection threshold (noise threshold units: NTU). For a given noise level, the stimuli were presented in random order and all target-flanker separations were mixed (‘Mix’ procedure). Each block consisted of 20 trials at each of the four target-flanker separations (80 trials per block). There were four blocks, each with a different external noise level (0, 0.5, 1, and 2 NTU). The starting noise level was randomized between participants. Participants repeated each noise level three times for a total of 960 trials (20 × 4 flank distances X 4 noise levels X 3 repetitions).

There were also a pilot experiment employing a set of 6 distances as in Polat and Sagi (2007). In this experiment target contrast was increased with increasing noise level to yield (approximately) constant sensitivity.

### Signal detection theory (SDT) analysis

We used the standard definitions of the sensitivity (*d*’) and the internal criterion (*c*) (Green & Swets, 1966):

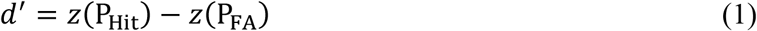

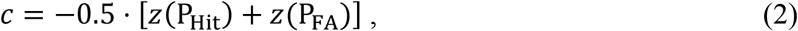

where P_Hit_ is the probability that an observer correctly reported that the target is present in target-present trials, P_FA_ is the probability that the observer incorrectly reported that the target is present in target-absent trials, and *z* is the inverse cumulative normal distribution function. To avoid saturation, the P_Hit_ and P_FA_ probabilities were clipped to the range 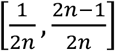, where *n* is the number of trials in the measurement.

### Reaction time (RT) behavioral analysis

We binned trials by categorizing RT into four equal-quantity bins from the fastest to the slowest. Binning was done separately for each experimental block and each trial type to avoid confounds (observer × experimental block × target-flanker distance × target stimulus (present/absent)). Task performance was quantified in each RT bin using the standard measures of sensitivity (*d*’, Eq. 1) and decision criterion (*c*, Eq. 2) from signal detection theory (SDT) (Green & Swets, 1966).

### Statistics

All statistics were assessed using linear mixed-effects models (fitlme(), Matlab2013a, The MathWorks, Inc., Natick, Massachusetts, United States), as detailed in the text. We consider two main factors contributing to criterion modulation, target-flanker distance (D_tf_) and RT. We tested for significant contributions of these two factors and their interactions to the measured criterion (c, Eq. 2), assuming:

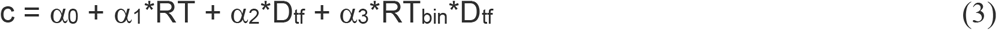

with D_tf_ (6 or 4 levels, see Table 1, “Target-flanker distance”) and RT_bin_ (4 bins, 0:3, fast to slow) defined as continuous effect, and observers as random effects (slopes and intercepts). Our main interest is in in α_2_ that measures the change in c as a result of increasing D_tf_, and α_3_ that measures the RT dependent addition to α_1,_ so that the D_tf_ slope equals α_2_+α_3*_RT_bin_. This simple model accounted for much of the variance in the data, with adjusted *R*^*2*^ between 0.5 and 0.8, where the lower values were obtained in the external-noise experiments, and in experiments with patients.

## Results

First, we confirm that, for the dataset analyzed here, the criterion dependency on distance differs between the two experimental conditions ‘Fix’ and ‘Mix’. The original results are presented in Figure 2B. In the ‘Fix’ condition, observers better adjust their criterion, as seen in the reduced dependency on target-flanker distance (smaller c(D_tf_) slope), a consequence of the different decision requirements presumably imposed by the mixing of different target-flanker distances in the ‘Mix’ condition. Indeed, testing the dataset analyzed here with the linear mixed-effects model for effect of experimental condition (‘Mix’: E_mix_ = 1, ‘Fix’: E_fix_ = 0), assuming c = α_0_ + α_1_*E_mix_ + α_2_*D_tf_ + α_3_*E_mix_*D_tf_, shows a significant effect of condition on the c(D_tf_) slope (α_3_=.035, *t*_(260)_ = 2.97, *p* =.003), with the slope more than doubled in the Mixed condition (α_2_+α_3_=0.061 vs α_2_=026). The c(D_tf_) slope in the ‘Fix’ condition (α_2_= 0.026) did not reach statistical significance (p=.09). The intercept showed no statistical difference between conditions (α_1_ =−0.19, p=.09).

### P1: Effects of RT on the criterion slope

To test our first prediction (P1), we examine the dependence of the c(Dtf) slope on RT. First, we consider the experimental data of the ‘Mix’ condition from Polat and Sagi (2007). In this experiment, trials having different target-flanker distances were mixed within a block (Figure 2A). Our RT analysis, presented in Figure 4A, clearly show that the criterion (*c*) had lower values for slower RTs, especially at larger target-flanker distances (D_tf_), implying that the criterion slope as a function of distance is reduced with increasing RT. This claim is strongly supported by the linear mixed effects model described in Eq. 1, showing the interaction between the RT bin index and distance (α_3_= −0.02) to be significant (*t*_(140)_ = −3.67, *p* <.001). The c(D_tf_) slope of the fastest RT bin (α_2_= 0.09) was significant (*t*_(140)_ = 4.68, *p* <.001), while the slope at the slowest RT bin (α_2_+3*α_3_= 0.03) touched statistical significance (*t*_(140)_ = 1.97, *p*=.05), suggesting criterion modulation at slow RTs.

**Figure 4:**
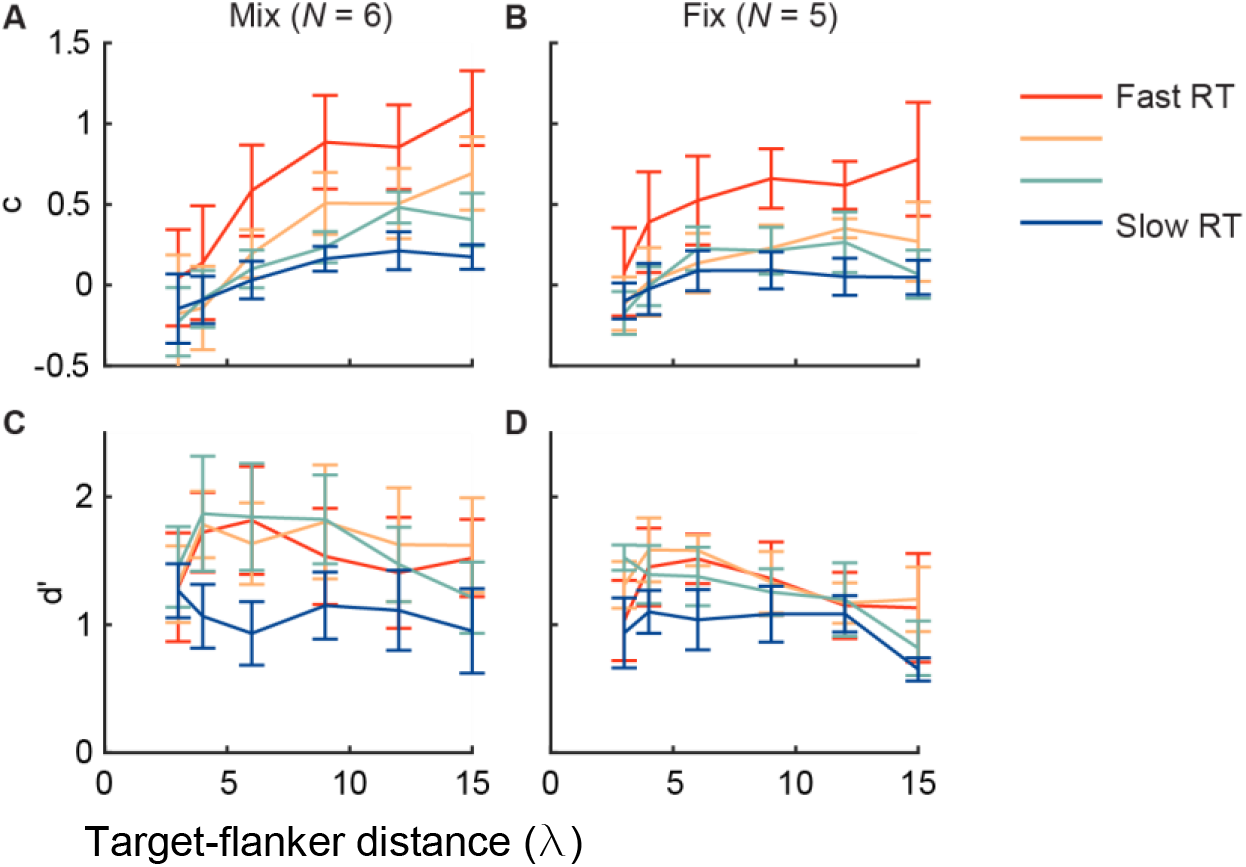
Target-flanker distance and RT. Analysis of (Polat & Sagi, 2007). (**A and C**) Shown are (**A**) criterion (c, Eq. 2) and (**C**) sensitivity (d’, Eq. 1), as a function of the target-flanker distance (in units of wavelength, λ), when the different target-flanker distances are mixed within blocks (“Mix” experiment). Data were split into four bins (colors) by sorting the measured RTs. The results indicated: (i) a more biased criterion setting in the faster RTs, (ii) a weaker modulation of the criterion by the flankers in the slower RTs, and (iii) a flanker-dependent criterion in all RTs, even the slowest. The sensitivity (d’) was lower with slower RTs, but only in the slowest time bin (blue). (**B and D**) Behavioral data of the “Fix” experiment, whereby trials are blocked by the target-flanker distance, showing reduced effects. Error bars are ±1 SEM.

We note here that criterion modulation is different from sensitivity modulation. Polat and Sagi (2007) found a small but significant modulation of sensitivity (*d*’) by the flankers, as seen in Figure 2C for the available data. (Note that a much stronger modulation of sensitivity by the flankers is found when the detection task is replaced by a 2AFC discrimination task (Polat & Sagi, 1993, 2007).) The expected sensitivity modulation with distance is non-monotonic, showing maximal effect at a distance of 3λ. Applying the RT analysis here revealed a gradual reduction of *d*’ in slower RTs, with values decreasing from ~1.5 in fast RTs, to ~1 in slow RTs (Figure 4C; *t*_(142)_ = −4.01, *p* <.001 for modulation of *d*’ by RT when ignoring the target-flanker distance). There was no significant interaction between RT and the target-flanker distance (*t*_(140)_ = −0.72, *p*=.5). Importantly, criterion and sensitivity exhibited different modulations by RT: the criterion was modulated to a much greater extent than sensitivity, and its modulation dynamics differed (Figure 4A vs. Figure 4C).

RT effects in the ‘Fix’ condition are qualitatively similar to the effects observed in the ‘Mix’ condition (Figure 4A vs 4B). Here, the linear mixed effects model described in Eq 1, showed the interaction between the RT bin index and distance (α_3_= −0.01) to be significant (*t*_(116)_ = −2.86, *p*=.005). The c(D_tf_) slope of the fastest RT bin (α_2_= 0.04) was significant (*t*_(116)_ = 3.07, *p*=.003), while the slope at the slowest RT bin (α_1_+3*α_3_= 0.01) was not statistically different from zero (*t*_(116)_ = 0.64, *p*=.52), suggesting, as predicted, no criterion modulation at slow RTs when target-flankers distances are blocked.

### P2: Effects of added external noise on the interaction between RT and c(Dtf)

Next, we consider experiments whereby different levels of external noise are added to the target (see the Methods, and Zomet et al. (2016)) using a procedure that is otherwise identical to the one considered in the previous section. Different noise levels were added in different blocks, randomly permuted between observers (see the Methods, Zomet et al. (2016)). We consider two variants: one where contrast was maintained when noise was increased, leading to reduced *d*’ when the noise level increased (“Main”, *N* = 12 observers, Figure A5). The other, an unpublished pilot of that study, whereby the target contrast was increased with noise to maintain a roughly fixed *d*’ (“Pilot”, *N* = 7, Figure A6; this pilot is particularly convenient for the purpose of the RT analyses performed here by virtue of having a large number of trials per observer, see Table 1).

First, we consider the case when the level of external noise was zero (“No noise”, Figures 4BC, A3, A4). In this condition, the experiment is equivalent to the one considered in the previous section (‘Mix’ condition), so we expect similar results. The results indeed show a strong interaction between RT and the c(Dtf) slope. Specifically, the c(Dtf) slope was reduced with RT (“Pilot”: *t*_(164)_ = −4.66, *p* <.001; “Main”: *t*_(188)_ = −3.01, *p*=.003). In addition, the criterion slope at the slowest RT bin was positive (“Pilot”: *t*_(164)_ = 4.62, *p* <.001; “Main”: *t*_(188)_ = −3.88, *p* <.001), as predicted for the ‘Mix’ condition (the estimated slopes are presented in Figure 8).

To test prediction P2, we examine the interaction between RT and c(D_tf_) slope in the presence of noise. As shown in Figure 5B and Figure 8, introducing noise reduces the interaction of criterion and RT. This is seen in Figure 8 as more similar c(Dtf) slopes for the fast and slow RT bins in the presence of noise. There is one exception, when noise level is 0.75*noise-threshold in the pilot experiment (*t*_(164)_ = −3.39, *p*<0.001), otherwise, as predicted, none of the interactions reach statistical significance (*p=0*.*1-0*.*9*). This effect is not due to a reduction in sensitivity (*d*’), as shown in Figure 5C, where the target contrast was increased with the added noise, so that sensitivity was fixed.

**Figure 5:**
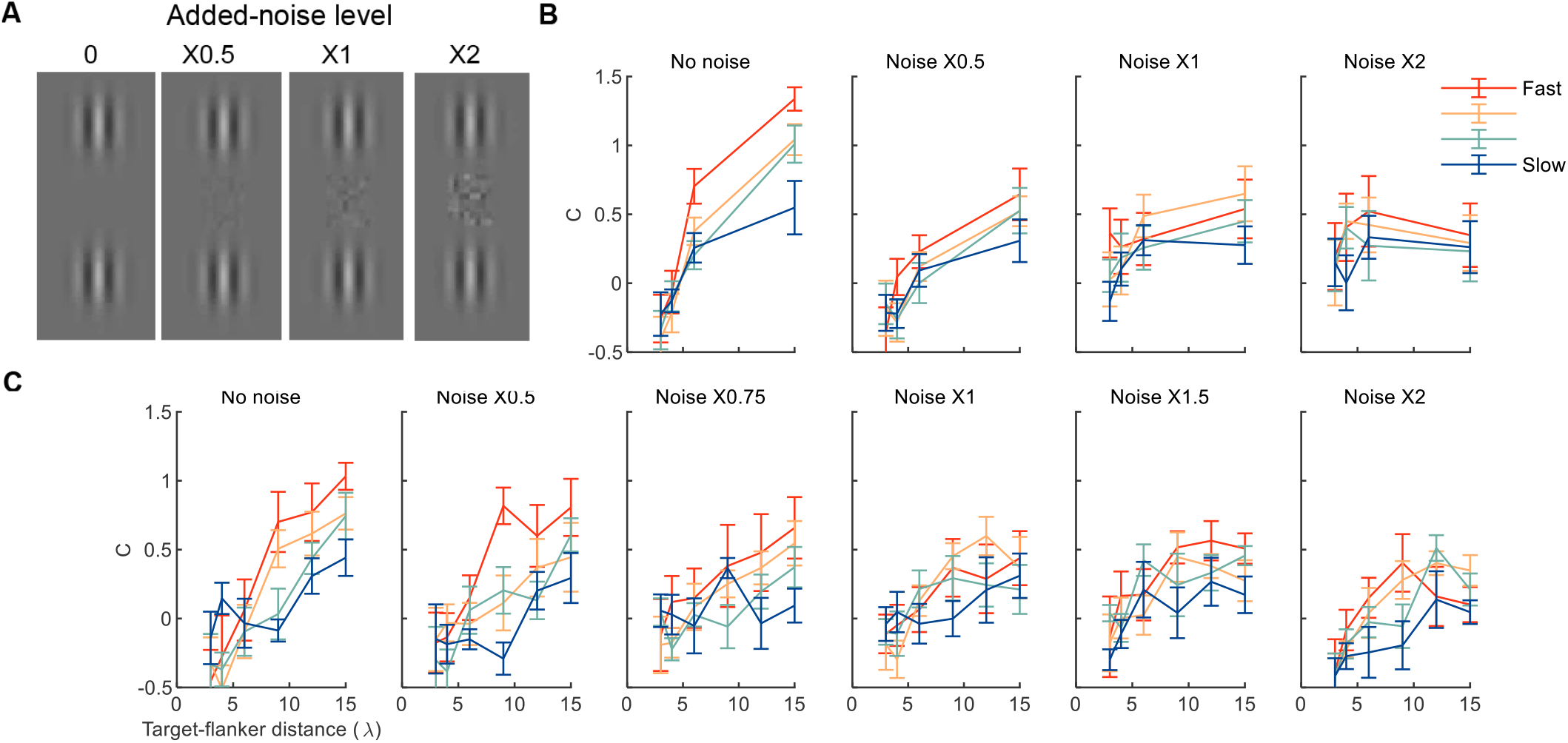
External noise and the RT effect. Analysis of Zomet et al. (2016). (**A**) Example stimuli with different added noise levels. (**B**) The “main” experiment where added noise reduces sensitivity (d’). (**C**) The “pilot” experiment where target contrast is increased with noise, so that sensitivity remains stable across noise levels. The error bars are ±1 SEM.

### P3: Effects of added external noise on the c(Dtf) slope

Of interest here are the results with the highest noise level. Here we expect the “Main” experiment and the “Pilot” to diverge. Indeed, examining the c(Dtf) slopes at the slowest RT, show it to be significant in the “Pilot” experiment, where target contrast was increased with noise (*t*_(164)_ = −3.01, *p*=.003), but not in the “Main” experiments where target contrast was not scaled with noise (*t*_(172)_ = 0.51, *p*=.5).

### P4: Individuals with slow RTs show no RT effect for criterion

Next, we applied the RT analysis to the experimental data of Zomet et al. (2008). In this study, patients diagnosed with depression (“Patients”) and their matched controls (“Controls”) performed an experiment similar to the ones analyzed above. Specifically, different target-flanker distances were mixed within a block, and there was no noise, comparable to the ‘Mix’ experiment of Polat and Sagi (2007) and the no-noise condition of Zomet et al. (2016). Unlike these studies, the number of trials was lower, and observers were, on average, older and slower (see Table 1).

The results of the RT analysis are presented in Figure 6. The criterion slope as a function of the target-flanker distance was reduced with RT, though not significantly for the patients (RT:slope interaction, “Controls”: *t*_(764)_ = −2.41, *p*=.02; “Patients”: t_(644)_ = −0.52, *p* =.6). The sensitivity (d’) was reduced with RT (“Controls”: *t*_(764)_ = −2.22, *p* =0.03; “Patients”: *t*_(644)_ = −3.06, *p* =.002), independent of the target-flanker distance (“Controls”: *t*_(764)_ = 0.19, *p*=.85; “Patients”: *t*_(644)_ = −1.46, *p*=.14).

**Figure 6:**
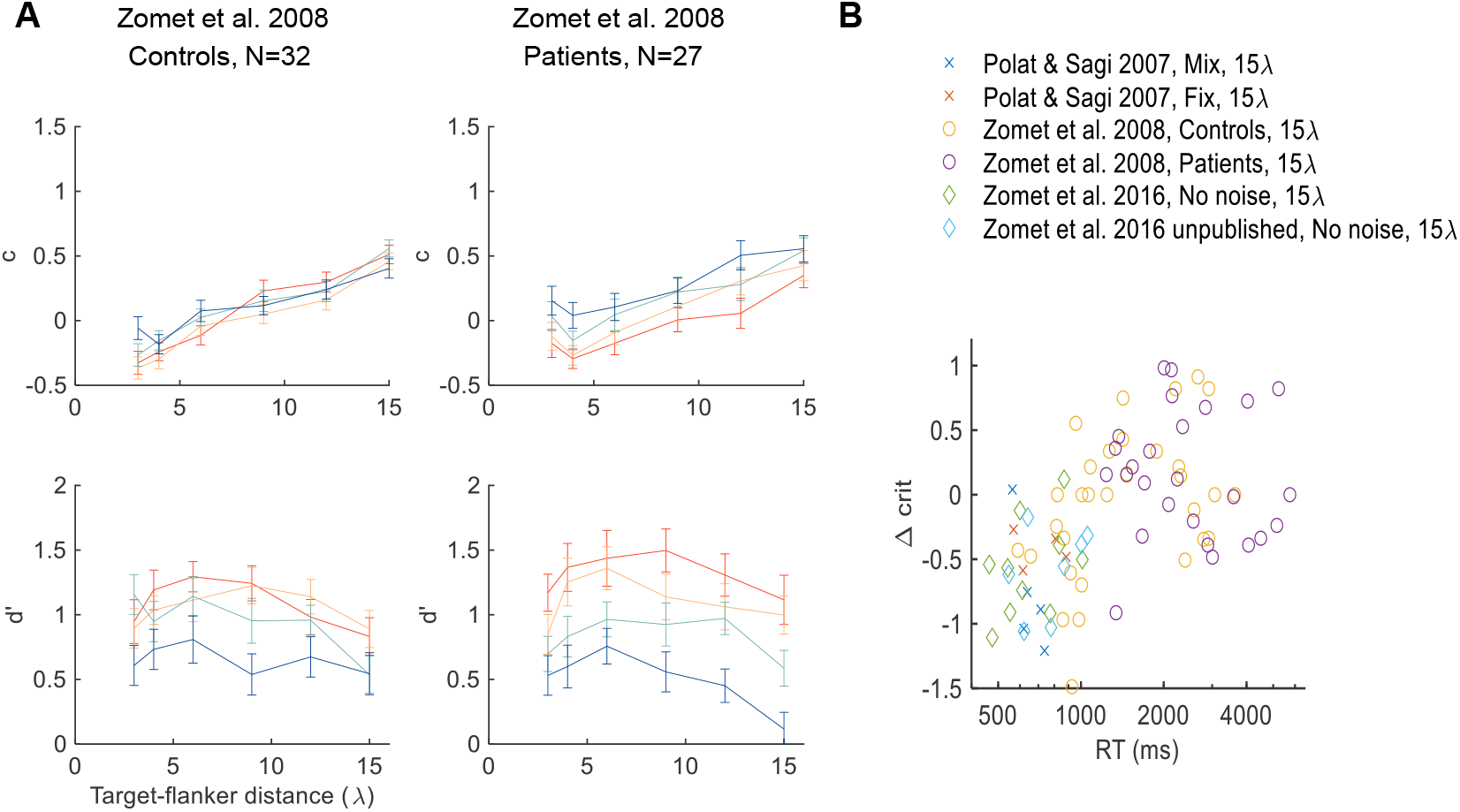
The RT effect in slow individuals. **(A)** RT analysis of Zomet et al. (2008). Annotations follow Figure 4. The results showed no RT effect for criterion in depressive patients and controls. (**B**) Shown for each observer is the change in criterion (Δc) from the fastest to the slowest RT bin of the 15λ measurement, as a function of the mean RT. The results indicated a correlation between RT and Δc between experiments, and within experiments for the Zomet et al. (2008) control data. The procedural differences between experiments are summarized in Table 1.

To explain the small effect of RT on criterion, we considered the simple idea described in Dekel and Sagi (2020a): that individuals with reduced bias can be explained by having slower decision times. Indeed, we found that observers in Zomet et al. (2008) had extremely slow RTs, showing, on average, ~1600 ms for Controls, and ~2700 ms for Patients (see the x-axis of Figure 6B). This is much slower than the ~650 ms measured for observers in similar experimental conditions (Figures 4A and 5 “No noise”). Overall, the differences between experiments can possibly be explained by RT. In addition, we considered differences between observers in the “Control” group of Zomet et al. (2008), where wide inter-individual differences in RT were measured (see the x-axis of Figure 6B). We correlated individual RT with the individual size of criterion modulation by RT (for 15λ distance), finding a significant correlation (adjusted *R*=0.54, *p*<.001; Figure 6B). To summarize, the difference between the experiment by Zomet et al. (2008) and the other experiments appears to be well explained by slower RTs. In addition, the difference in RT between experiments and observers can possibly be attributed to age and/or practice effects (Table 1).

### Summary of results

The experimental results show a clear dependency of decision bias on the target-flanker distance, which is expected to disappear at slower response times if caused only by criterion setting (*sp* in *DDM*), but not if caused by changes in the internal distributions associated with the different stimuli (affecting drift rate in *DDM*). Our results, summarized in Figures 7 and 8, support the latter.

**Figure 7:**
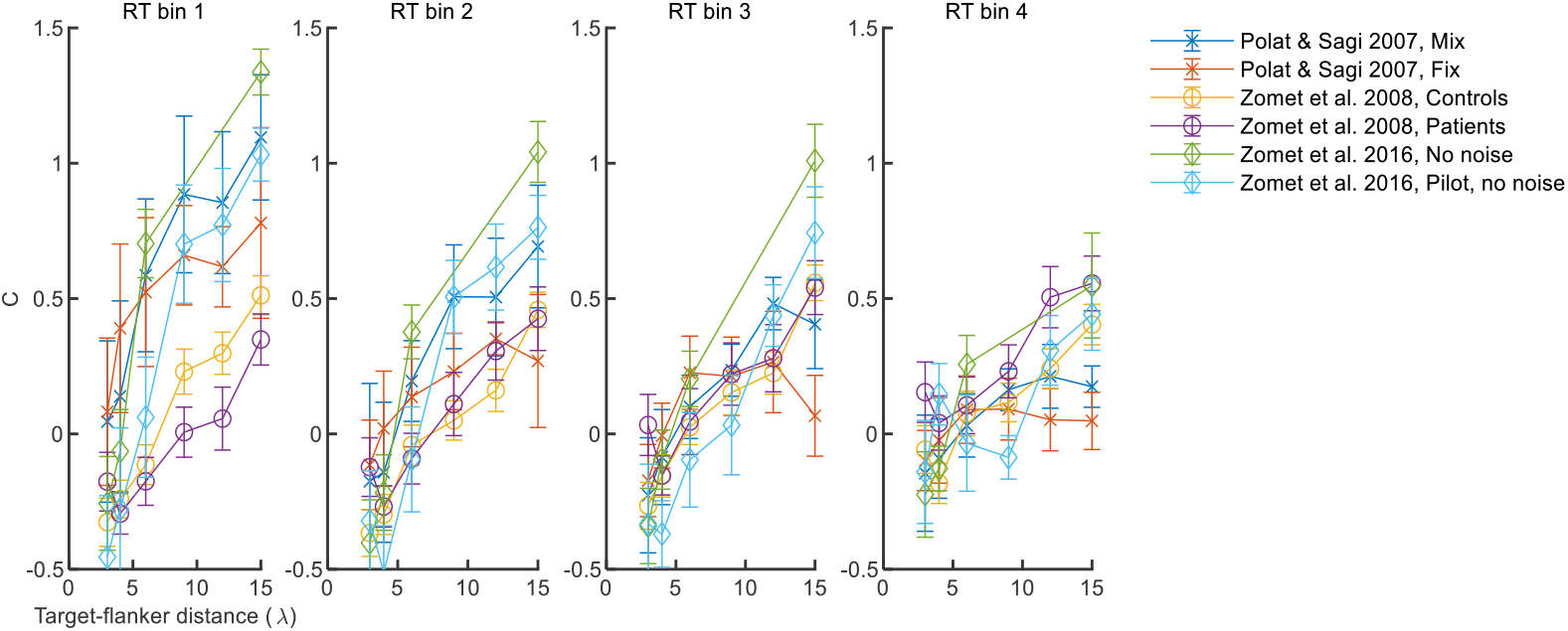
The differences between experiments disappear with slow RTs. Shown for all experiments without noise are the criterion measurements as a function of the target-flanker distance in four RT bins. It can be observed that the experimental differences are abolished in the slowest RT bin (bin 4).

Figure 7 depicts c(D_tf_) functions for the four RT bins (1 being the fastest). In the slowest RT bin the differences between experiments and conditions are largely abolished (but note the flat ‘Fix’ curve), as clearly seen when comparing bin 4 (slow RTs) with bin 1 (fast RTs). Figure 8 depicts the c(D_tf_) slopes for the fastest and slowest RT bins, for all the experiments analyzed here, fitted to data using Eq. 3.

### DDM predictions

To try to understand the observed interactions with RT, we considered the idea that the detection task is performed by gradual accumulation of noisy evidence, as in the standard drift diffusion model (DDM) (Gold & Shadlen, 2001; Ratcliff et al., 2016). Here we considered some general effects of model parameters on its behavior (more detailed model parametrization and fitting results are provided in the Appendix, Figure A2). As described in the Introduction, two important parameters in the model affect the criterion: (i) the starting point (*sp*) of the accumulation process, which leads to increased bias in faster decisions if put closer to one of the bounds (Figure 9, second row) (Dekel & Sagi, 2020b), and (ii) the rates at which evidence is accumulated for target-present and target-absent trials (Figure 9, first row). As shown in Figure 9, this simple idea can explain the criterion measurements quite well. We provide here some intuition as to how this model works by describing the predictions of four simple model variations and analyzing the effect of the drift rate and the starting point on the results. The analysis is sufficiently general for its predictions to be independent of the specific assumptions made on the drift rate (see Introduction). For simplicity, drift rate is considered here as an abstract variable monotonically related to the log likelihood ratio *(LLR)* (Gold & Shadlen, 2001).

**Figure 8:**
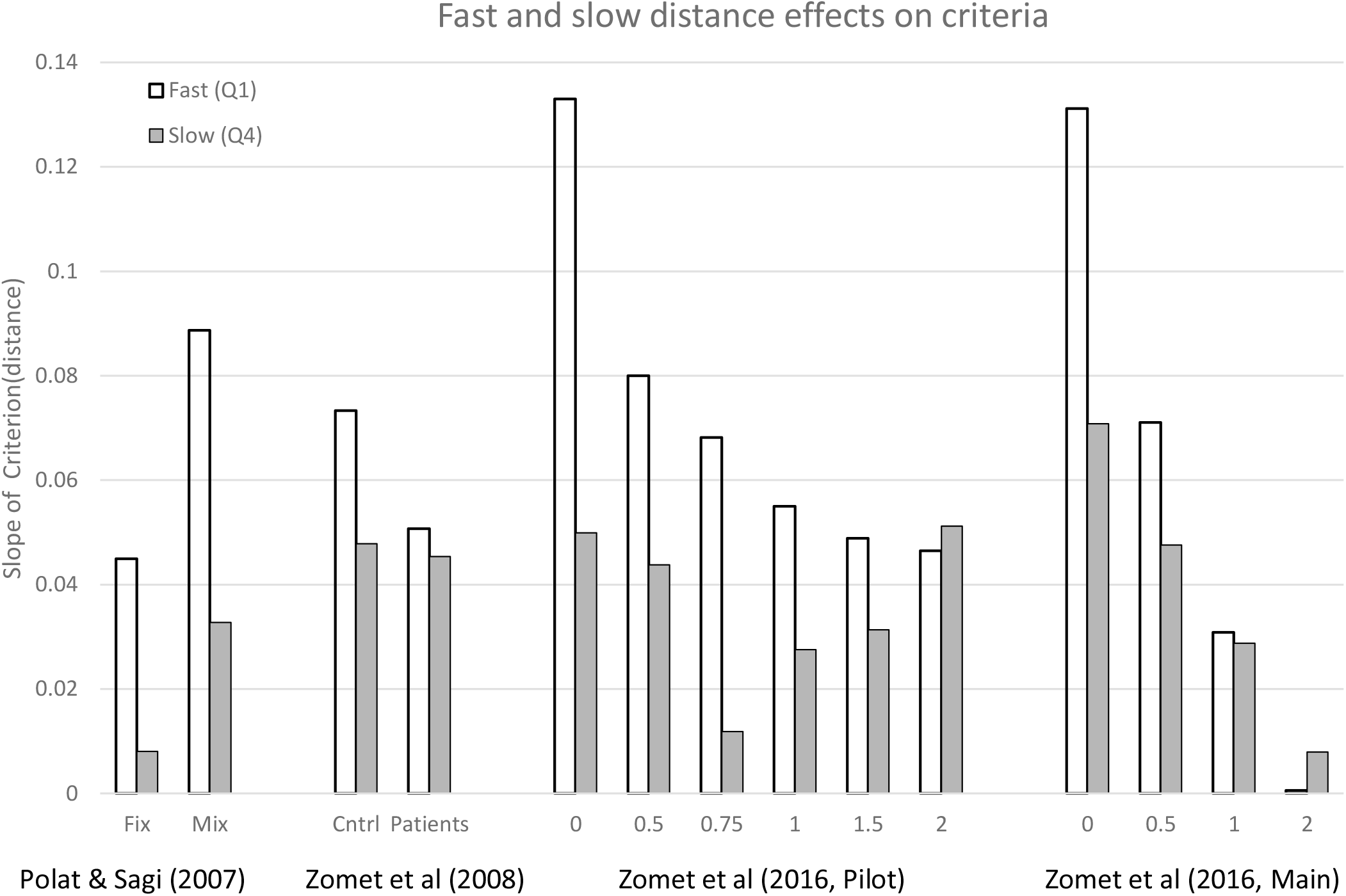
Fast (Q1: 1^st^ RT quantile) and slow (Q4: 4^th^ RT quantile) distance effects on criteria: a summary of all studies. Shown are the c(Dtf) slopes fitted to the respective RT bins using the linear model described by Eq. 3. All fast distance effects, but one (highest noise level in Zomet 2016 Main), are statistically significant (see the Results and Figures A5 and A6). The slow effects are more uniform across conditions, all significant except for the Fix condition (see P1) and the noise-amp=0.75 condition in the “Pilot” experiment (see Discussion), and the noise-amp=2 condition in the “Main” experiment (see P3).

**Figure 9:**
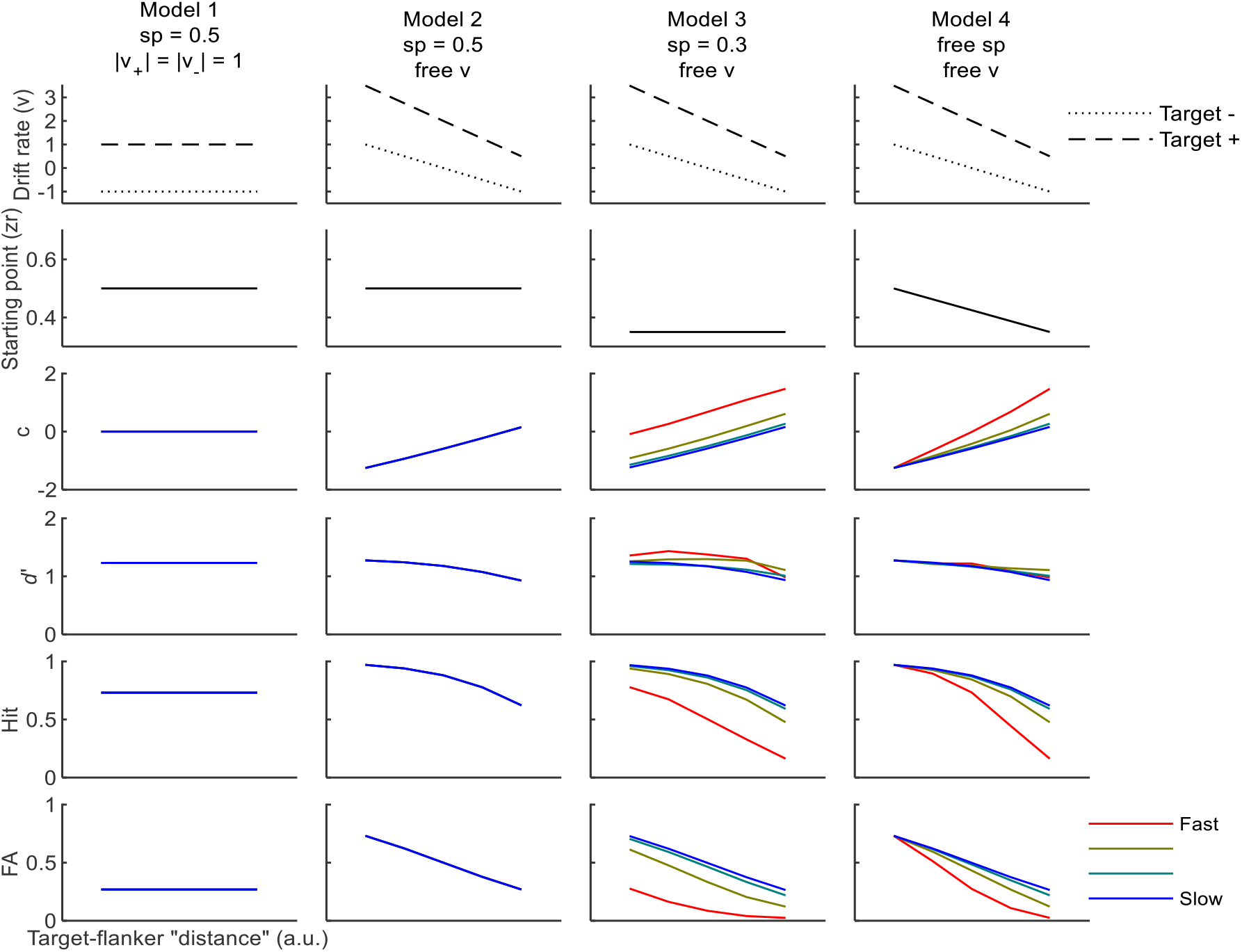
Modeling the influence of flankers on target detection using DDM. Shown are the parameters (the first two rows) and the theoretical predictions (the last four rows) of the four models described in the text. The sensitivity estimate (d’, Eq. 1) is approximately the difference in drift rate between the target present and absent trials. In models 3 and 4, the starting point is biased (sp<0.5), leading to an overall reduction in criterion (c, Eq. 2) with RT. In Model 4, the reduction in slope with RT can be explained by a flanker-dependent starting point (variable sp). Overall, Model 4 manages to explain both the reduction in overall criterion with RT, and the reduction in c(distance) slope with RT. See the text for model details and Figure A2 for fitting Model 4 to experimental data.

#### Model 1: An unbiased flanker-independent starting point, target-dependent drift rate

Here, the drift rate (*v*) depends on the target stimulus (*v*_*+*_=+1 for target present, *v_−_*=−1 for target absent), and the starting point (*sp*) is fixed at 0.5 (unbiased). This model predicts no bias (*c*=0) at all RTs, with fixed *d*’, independent of distance. This model fails to explain the data.

#### Model 2: As in Model 1 with a flanker-dependent drift rate

Here, at short distances, the difference between *v*_*+*_ and *v*_*-*_ is increased, thus showing larger *d*’ values at these distances. Both *v*_*+*_ and *v*_*-*_ are positive at short distances (supporting positive responses), thus showing a negative criterion (*c*<0) at these distances, and RT independent. This model can explain the distance dependence, but not the RT dependence, of the measure criterion,

#### Model 3: A biased flanker-independent starting point (*sp*), flanker-dependent drift rate

The same as in Model 2, but with a biased starting point (*sp*). The effect of this bias (*sp*<0.5) is short term, resulting in higher criteria (*c*) in trials with fast RTs, and with slow RTs showing criteria equal to those produced by Model 2. The RT effect on *c* is distance independent. This model predicts an RT effect but cannot explain the dependence of the criterion slope on RT (Figure 8).

#### Model 4: A flanker-dependent starting point (*sp*) and drift rate

Like Model 3 but with *sp* allowed to change with distance, resulting in distance-dependent RT effects (RT-dependent slope of the *c(distance)* curve). This model captures the main features of the experimental results.

Overall, the DDM framework considered here can qualitatively account for the most salient RT effects, but a precise modeling, requiring several more parameters and assumptions, remains somewhat speculative. In the Appendix (Figure A2) we present a detailed fit of Model 4 to the behavioral data, using the Fast-DM software with the Kolmogorov-Smirnov (KS) setting (Voss & Voss, 2007)..

### DDM drift rate and SDT *d*’

Note that the DDM analysis allows for the derivation of target-flanker distance-dependent sensitivity functions for both target-present and target-absent conditions (the idea in Figure 9, fitting in Figure A2), unlike the previously used SDT methods that provided only a differential sensitivity estimate (i.e., *d*’). As seen in Figure A2, the fitted drift rates for both target-present and target-absent gradually decrease with longer target-flanker distances, as predicted by the theory illustrated in Figure 1. The differential sensitivity, which is the difference in the fitted drift rate between target-present and target-absent trials, was mostly fixed, showing a small decrease at longer target-flanker distances. This is consistent with the slightly improved *d*’ in the presence of proximal flankers. We noted that the fitted drift rate in the target-absent trials was usually negative. A negative drift indicates that the evidence supports the negative (target-absence) response. That is, the reduced sensory response in the target-absent trials is mapped into *negative values* (see the Discussion). This finding is reasonable in the sense that what is accumulated is target-presence evidence, with “zero” values corresponding to no evidence for or against target presence (see the Introduction).

## Discussion

Lateral masking affects the detection of low-contrast targets (Polat & Sagi, 1993). This effect is usually quantified using measures of behavior derived from signal detection theory (SDT) (Green & Swets, 1966): sensitivity (*d*’, Eq. (1)) and decision criterion (*c*, Eq. (2)). Although the bias-independent sensitivity measure (*d*’) is the standard measure used to quantify perceptual effects, our interest here lies in effects on decision criterion (bias), shown in Figure 2. SDT, with limited access to internal distributions (Gorea & Sagi, 2000), provides an observer-independent account of the biases found, as illustrated in Figure 3 (see the Introduction). Here, for the first time, we performed an RT analysis of the lateral masking effect, attempting to isolate the subjective (observer dependent) and objective (observer independent) factors underlying the observed biases in perception.

To explain the effects of RT, we extended the time-independent SDT explanation (Figure 1) by using the drift diffusion model (DDM). More specifically, we modeled the influence of the flankers on the detected target (Figure 1C: signal shift) as a change in the rate of evidence accumulation (drift rate), which leads to the time-independent effects on the measured criterion that are maintained even at the slowest RTs. The time-dependent effects on criterion were modeled as a change in the starting point (Figure 1B: criterion shift).

We found that mixing trials of different target-flankers distances affects mostly the slow RT trials. The fixed distance procedure, unlike the mixed distance procedure, produces negligible distance dependence at the slow RTs but not at the fast RTs (Results, P1). This result is explained by a mismatch between the actual internal distributions associated with the stimuli and the one (averaged across distances) used by the observer to make a decision concerning target presence.

We found that adding external noise (Zomet et al., 2016) affects mainly the fast RTs (Results, P2), equating fast and slow effects, implying that the influence of the starting point becomes negligible due to the presence of external noise when accumulation starts. For the condition where target contrast was not increased to compensate for the increased noise, we found reduced effects on criterion at both fast and slow RTs, as predicted by the change in the slope of the *LLR* function (P3, Figure 3).

In Zomet et al. (2008) the observers are slow (because of age or lack of practice); thus, we find the criterion effects to be much smaller, showing little or no dependence on RT (Results, P4). This agrees with our previous results showing that individual differences in TAE can be explained by RT differences (Dekel & Sagi, 2020a).

The presented RT analysis provides further insight into processes underlying statistical decisions. Of particular interest are limitations imposed on such decisions. We identified here a limit on the number of statistical distributions that can be efficiently learned by observers facing decisions on stimuli sampled from different distributions. Considering the example presented in Figure 3, limitations on the internal representations of response distributions are expected to be universally expressed, i.e., to introduce biases in slow responses. Indeed, slow distance-dependent biases are shown in 11 experimental conditions out of 14 (see Figure 8). As discussed above, we predicted two of the three exceptions. Of critical importance for the present theoretical framework is the reduced effect in the ‘Fix’ condition of Polat and Sagi (2007). Here, unlike in the ‘Mix’ condition, the different target-flanker distances are blocked, leaving us with only two internal distributions to learn (Signal and Noise); thus, the imposed constraints do not apply. A second predicted exception, described above (P3), concerns the absence of slow RT effects in the presence of high external noise. The third exception, not explained within the theoretical framework presented here, is the experimental condition with the external noise amplitude approaching noise threshold (noise amplitude=0.75threshold). We attribute this reduced distance effect to the increase in the False Alarm rate at slow RTs (otherwise present only at target locations near the flanker), and to the reduced sensitivity at all distances (d’ approaching zero), caused by external noise level close to threshold, as indicated by the results presented in Figure A5.

Within the context of the DDM framework, the slow RT effect can be quantified as drift rate asymmetry (Model 2-4, Figure 9, Figure A2), which is the sum of the up (target present) and down (target absent) drift rates (**ν**_**+**_**+ν**_**-**_). For an unbiased observer, this sum is expected to be zero. Figure 10A presents drift-rate asymmetry in the fitting results (Figure A2) for both the “Mix” and the “Fix” conditions of Polat and Sagi (2007), showing a marked difference between conditions. In contradistinction, biases due to shifts in the starting point (sp) are very similar in the ‘Mix’ and the ‘Fix’ conditions (Figure 10B). Accordingly, we attribute the differences in results between the ‘Mix’ and the ‘Fix’ conditions to objective factors affecting decisions, i.e., to differences in the internal responses, rather than to subjective factors such as priors (see the Introduction). We can conclude that the biases in the ‘Mix’ condition can be explained by the distance-dependent excitatory effects of the flankers, operating in both target present and absent trials, underlying the perceived filling-in.

**Figure 10:**
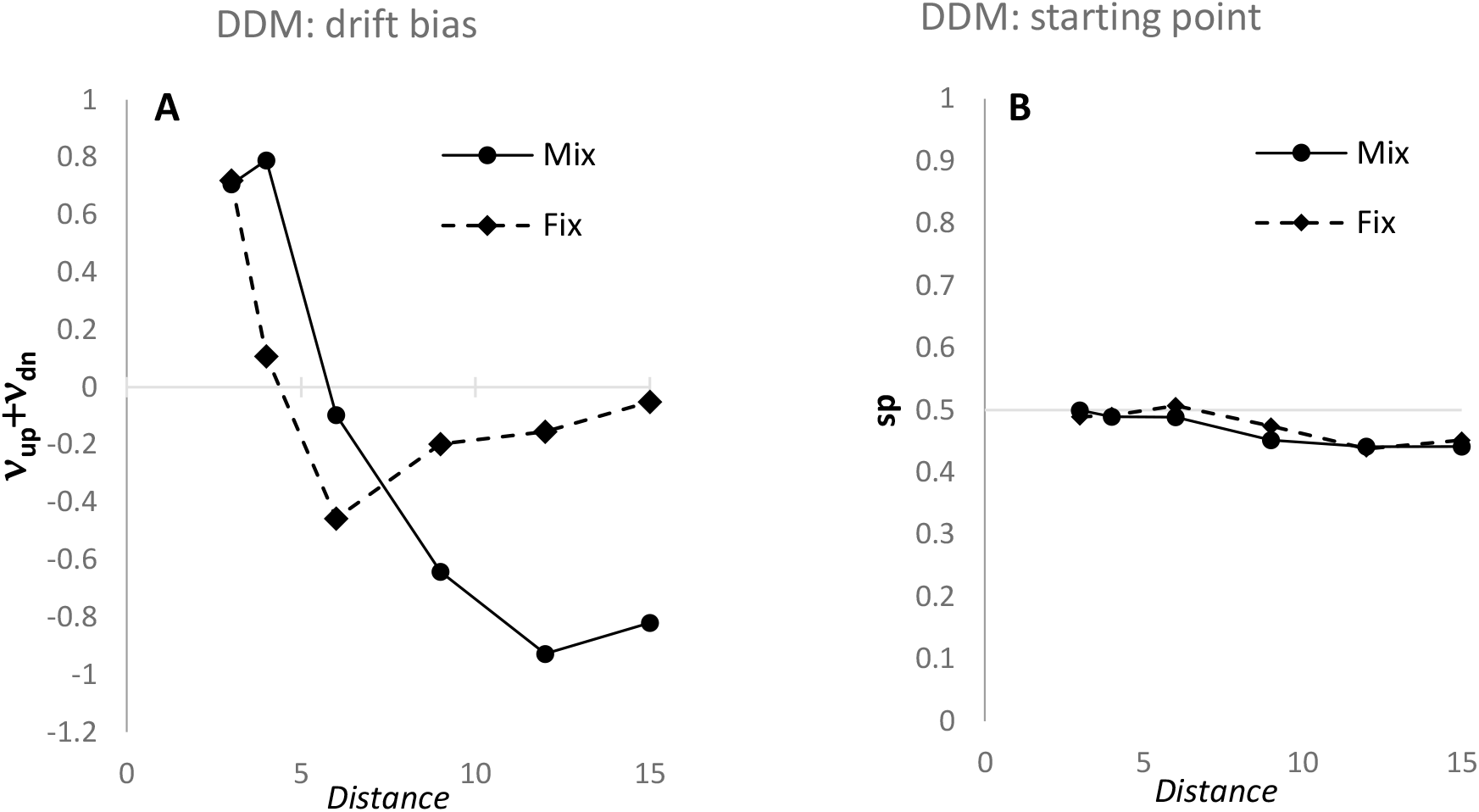
(**A**) Drift rate asymmetry, quantified as the sum of the up and down drifts, evaluated by fitting Model 4 (Figure A2) to the data of Polat and Sagi (2007). A positive bias leads to increased Hit and FA rates, and thus, to lower decision criteria. Note the difference between the two conditions. (**B**) Drift starting point (sp), evaluated by fitting Model 4 to the data of Polat and Sagi (2007). Biases due to prior information or payoff are indicated by deviations from sp=0.5. Both experimental conditions show unbiased sp at short distances but a lower sp at longer distances, resulting in lower Hit and FA rates at these distances (higher criteria).

Although additional modeling details remain somewhat speculative, the basic idea used here, of separately considering RT-dependent and RT-independent effects and interpreting them as changes in how evidence is interpreted (the drift rate) and what priors exist before the evidence (the starting point) seems quite robust.

## Acknowledgments

This research was supported by the Basic Research Foundation, administered by the Israel Academy of Science (grant # 2494/21, UP and DS).

### Appendix

**Figure A1:**
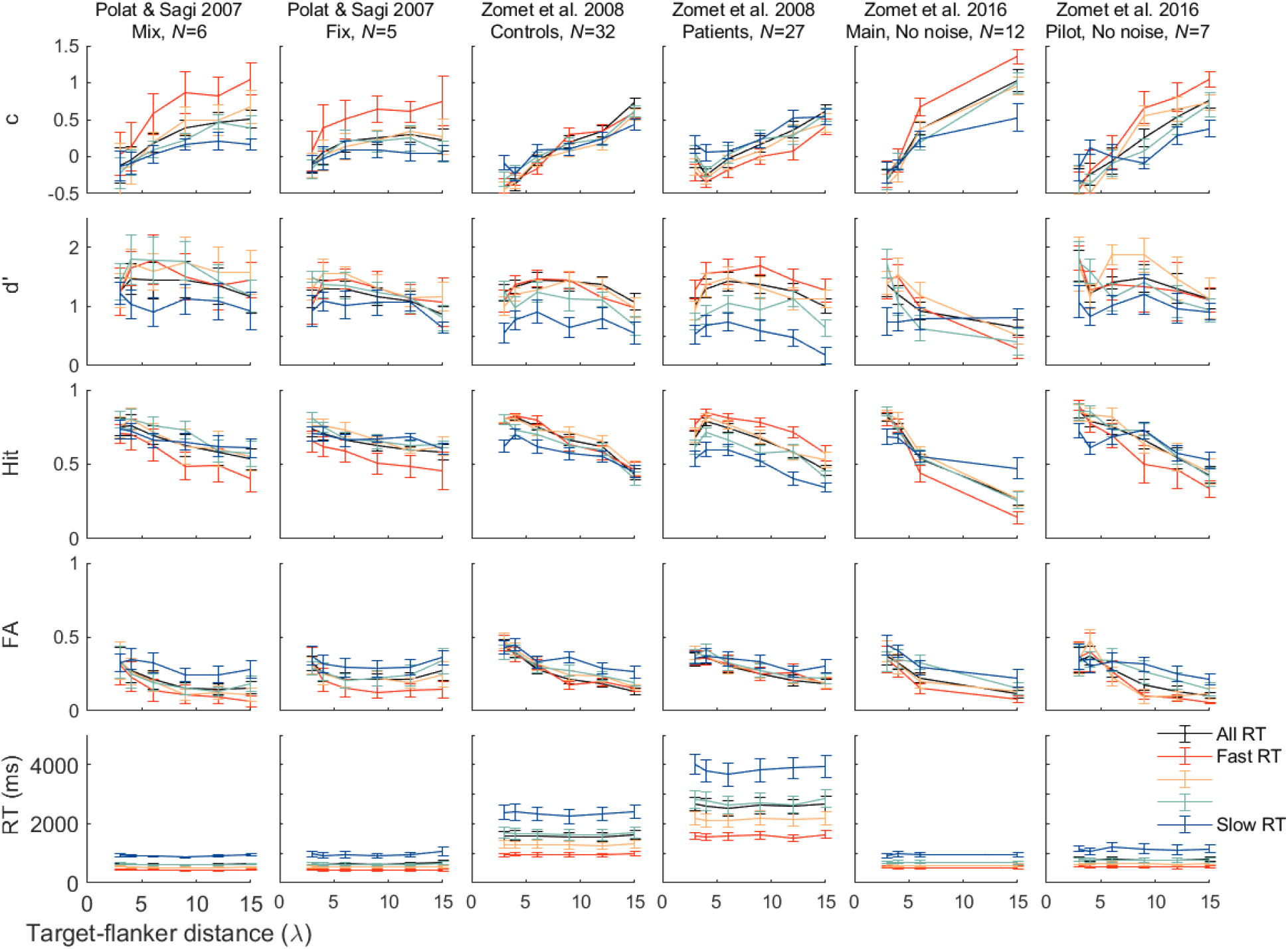
Target-flanker distance and RT. Shown, in six experiments (columns), are five behavioral measures (rows): criterion (c), sensitivity (d’), hit rate, false alarm rate, and RT, as a function of the target-flanker distance. Data were split into four bins (colors) by sorting the measured RTs. Error bars are ±1 SEM.

**Figure A2:**
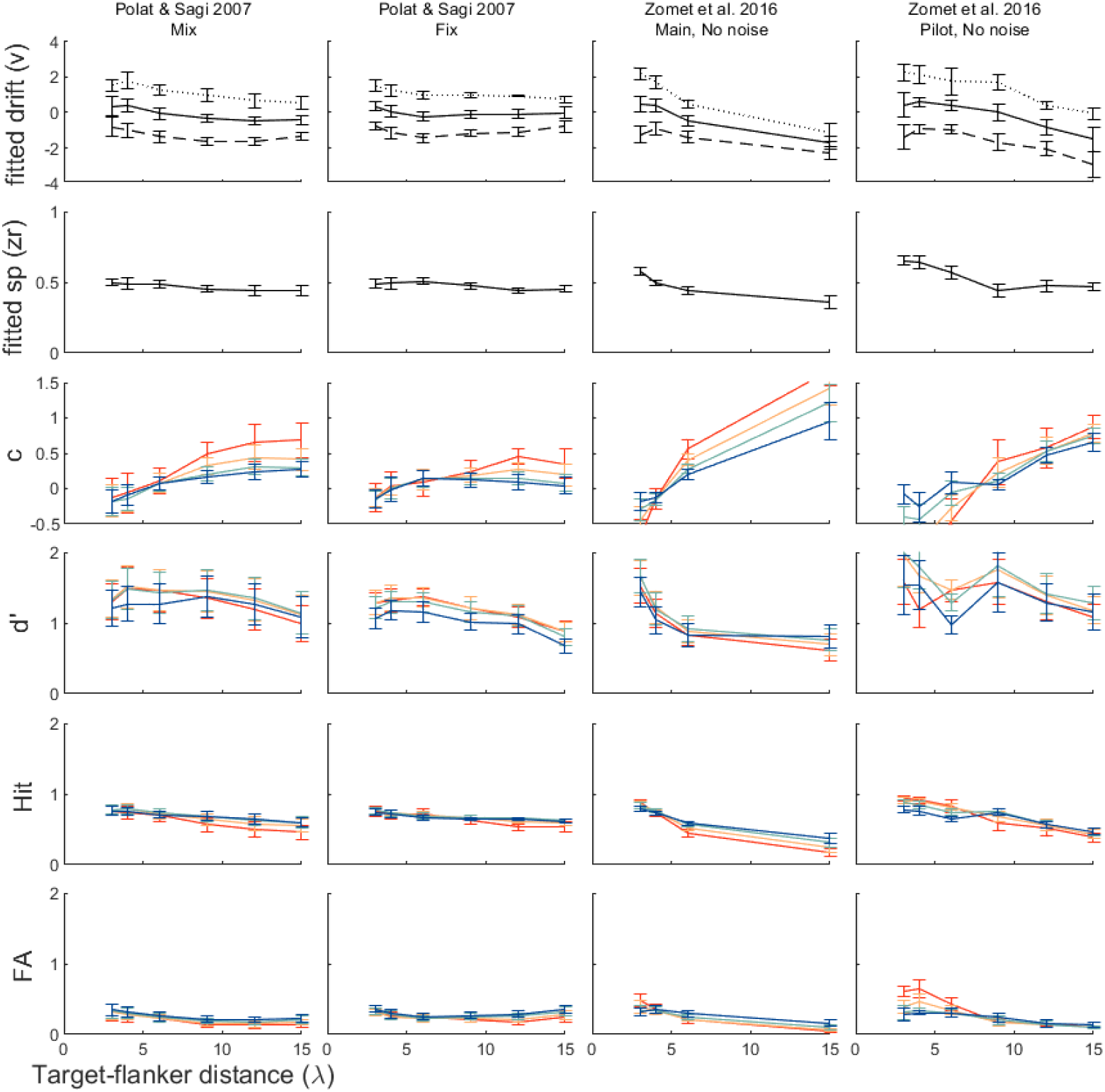
Fitting behavior to Model 4. Top two rows present the recovered parameters: drift rate (v_+_, v_-_ and their mean) in the first raw, and the starting points in the second raw, Rows 3-6 present the recovered behavior (c, d’, pHIT and pFA). To obtain analytical model distributions and to fit Model 4 to behavioral data, we used the Fast-DM software with the Kolmogorov-Smirnov (KS) setting (Voss & Voss, 2007). Fitting was performed separately for each observer. The drift rate (v) was fitted separately for each combination of the target stimulus (target present/absent) and the target-flanker distance (four or six possible distances) (8 or 12 parameters); the starting point (sp) was fitted separately for each target-flanker distance (four or six possible distances) (4 or 6 parameters), and the non-decision time (t_0_) and bounds-separation (a) were fitted once for all configurations (2 parameters). Overall, the model recovers the important features of the experimental results shown in Figure A1 (R^2^=0.78, comparing all 88 RT binned criteria, 3^rd^ raw, broken to experiments, from left to right, R^2^=0.87, 0.6, 0.93, 0.79).

**Figure A3:**
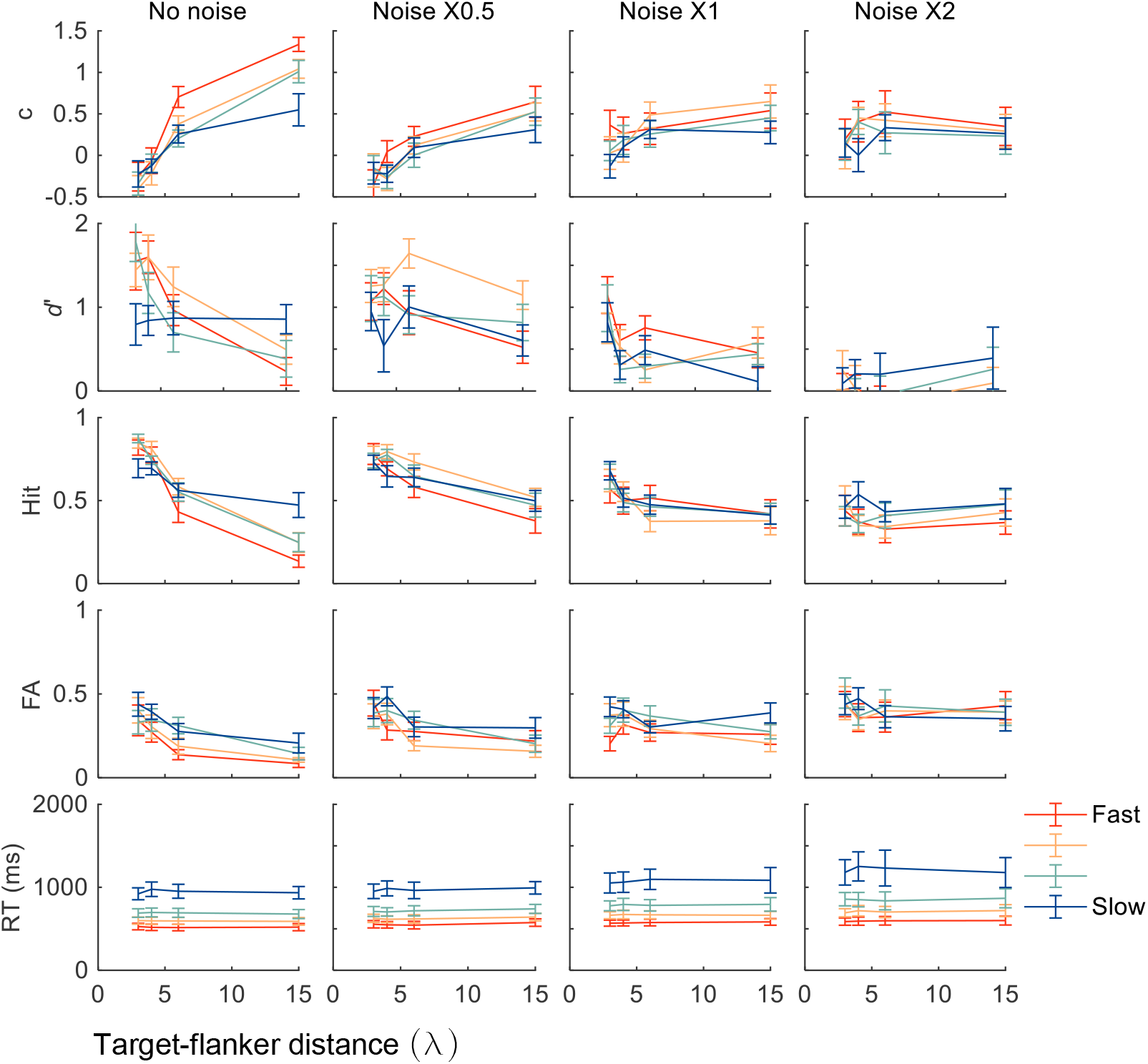
RT and external noise. RT analysis of the “main” experiment of (Zomet et al., 2016), and the layout following Figure A1. The first row is reproduced from Figure 5B.

**Figure A4:**
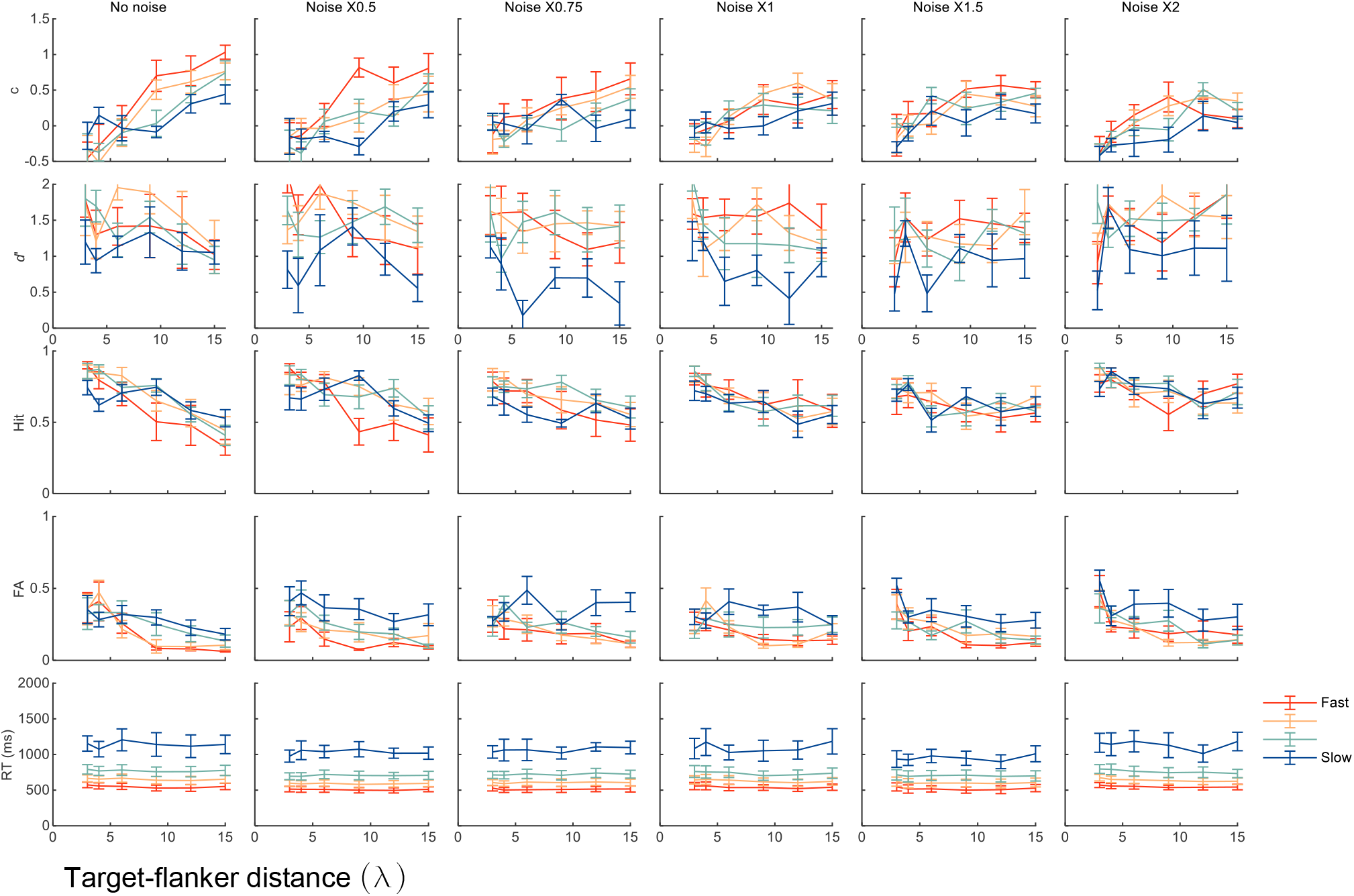
RT and external noise. RT analysis of the “pilot” experiment of (Zomet et al., 2016). The first row is reproduced from Figure 5C.

